# Self-organization of songbird neural sequences during social isolation

**DOI:** 10.1101/2022.02.18.480996

**Authors:** Emily L. Mackevicius, Shijie Gu, Natalia I. Denisenko, Michale S. Fee

**Affiliations:** McGovern Institute for Brain Research, Department of Brain and Cognitive Sciences, MIT

## Abstract

Behaviors emerge via a combination of experience and innate predis-positions. As the brain matures, it undergoes major changes in cellular, network and functional properties that can be due to sensory experience as well as developmental processes. In normal birdsong learning, neural sequences emerge to control song syllables learned from a tutor. Here, we disambiguate the role of experience and development in neural sequence formation by delaying exposure to a tutor. Using functional calcium imaging, we observe neural sequences in the absence of tutoring, demonstrating that experience is not necessary for the formation of sequences. However, after exposure to a tutor, pre-existing sequences can become tightly associated with new song syllables. Since we delayed tutoring, only half our birds learned new syllables following tutor exposure. The birds that failed to learn were the birds in which pre-tutoring neural sequences were most ‘crystallized’, that is, already tightly associated with their (untutored) song.

## 1 Introduction

On the one hand, sensory experience is known to be essential for the normal development of brain circuits. On the other hand, genetically specified developmental processes are also essential – we learn too quickly and from too sparse data to rely on sensory experience alone [1]. Thus, it appears that the brain is able to use genetically specified predispositions to fill in gaps in its sensory experience. When typical sensory experience is absent or delayed, certain aspects of brain development proceed anyway, while other aspects are delayed. This is true both in primary sensory systems [2, 3, 4, 5], and for more cognitive behaviors such as social interaction and language [6, 7, 8]. Brain circuits acquire structure and organization even in the absence of typical training inputs. Here we examine this self-organized structure, and what happens when sensory experience is reintroduced, in the context of songbird vocal learning.

Song learning is influenced by both auditory exposure to a particular tutor song, and by inherited preferences [9]. It is well known that songbirds, in the absence of exposure to a tutor bird, develop ‘isolate’ songs, with highly variable and atypical syllable rhythms [10, 11, 12]. However, when these ‘isolate’ songs are used as tutor songs, after two generations birds sing normally again, suggesting that an ‘innate’ preference filters what aspects of a tutor song are actually imitated [12]. Song imitation requires remarkably little total exposure to a tutor song – approximately 75 seconds total on a single day is enough for a bird to remember a song, and subsequently practice and imitate it [13]. Zebra finches, like many songbird species, are able to imitate songs of birds from other species, but when given a choice they prefer zebra finch song [14]. Furthermore, inherited genetic predispositions have a strong effect on both the precise tempo at which a zebra finch sings its song [15], as well as the particular learning styles of individual birds [16]. Thus, within the songbird brain we expect to see an interplay between developmentally specified and learned structure.

There are several possibilities for what happens in the brain during isolate song, and how it compares to typical (tutored) brain development. In typical birds, neurons in HVC are initially only weakly coupled to song, firing only at the onsets of syllables when birds are babbling subsong [17]. Then, as the song becomes more mature and repeatable, each HVC projection neuron fires at its own precise moment during the song, together forming a stable sequence of neural firing that tiles the song [17, 18, 19], in interplay with inhibitory neurons [20, 21]. This maturation process in HVC has been modeled as an initially random network of neurons that, with the right training inputs and plasticity rules, assembles into a chain of sequentially connected neurons [22, 23, 17] (Figure 1A). However, what happens in birds isolated from a tutor? Compared to typical adult zebra finch song, isolate song has a much less stable sequence of syllables and abnormally variable acoustic structure and timing [12]. In fact, aspects of isolate song resemble features of early babbling (subsong). Does HVC in isolate birds resemble that of subsong birds? Or does HVC mature to form sequences, even without experience of a tutor, and without the behavioral stereotypy seen in adult birds? We use functional calcium imaging in singing isolated birds to address these questions.

**Figure 1:**
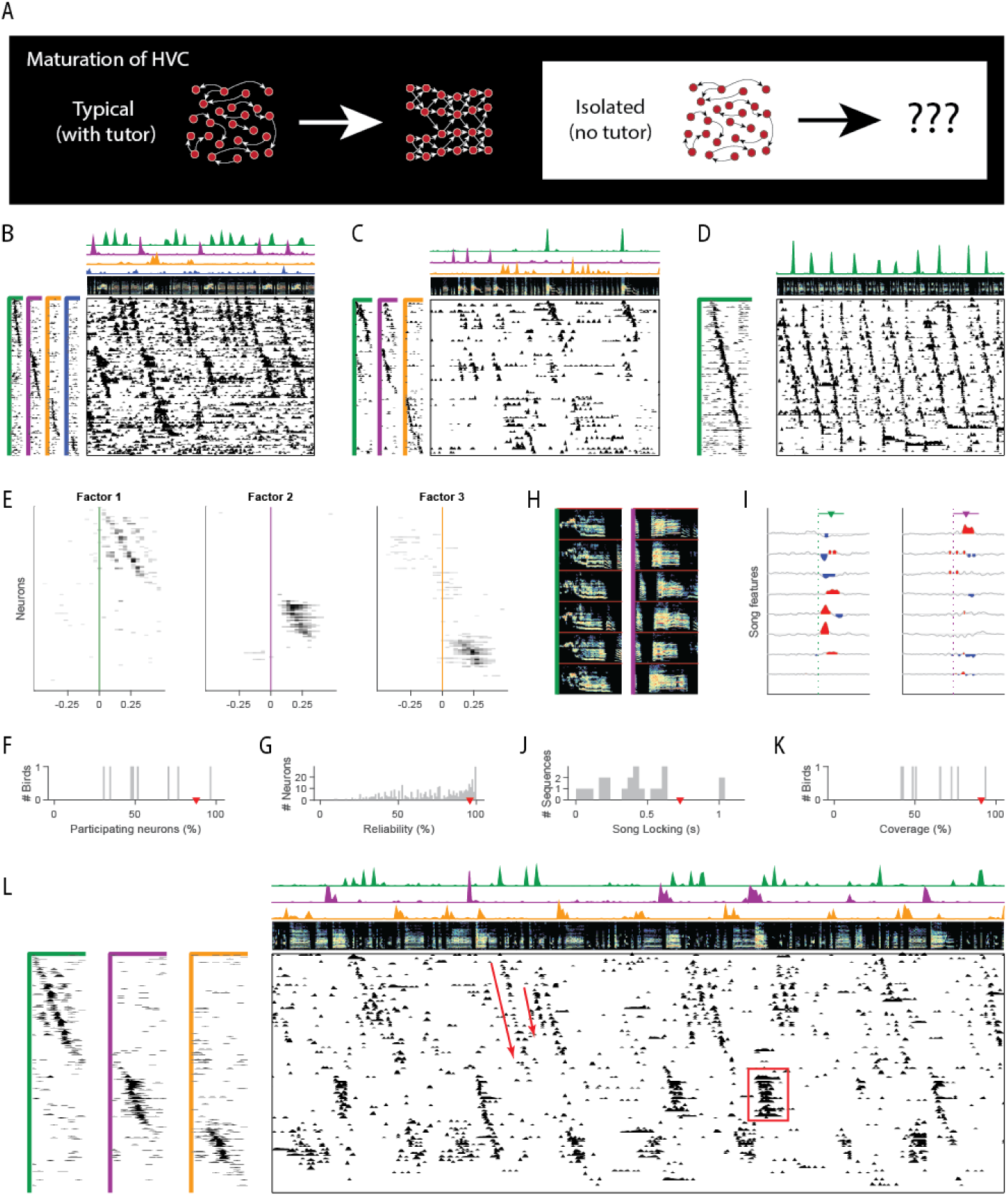
Sequences in isolated birds (A) Diagram of HVC maturation. In typically tutored birds, HVC sequences appear to grow and differentiate over time. (B) Example neural sequences recorded in a singing isolated bird (older juvenile, 61 dph). Main panel (lower right), functional calcium imaging recordings from 98 neurons for a duration of 6 s. Rows (neurons) sorted according to sequences (factors) extracted by unsupervised algorithm seqNMF (see Methods). (Above) Song spectrogram (0-10 kHz). The four sequence factor exemplars and timecourses are shown to the left and above, in corresponding colors. Duration of factor exemplars: 0.5 s. (C) Same as B, for another example isolated bird (adult, 117 dph). (D) Same as in B, for a typically tutored bird (adult, 217 dph). (E) Time-lagged cross correlation between each neuron and each of the three extracted factors recorded in a singing isolated bird (older juvenile, 68 dph). Only significant bins in the cross correlation are shown (p < 0.05, Bonferroni corrected, compared to a circularly-shifted control). (F, G, J, K) Sequence properties in isolated birds. For reference, median for typically tutored bird in D shown in red. (F) Percent of neurons participating in at least one extracted sequence. (G) Reliability of participating neurons across sequence renditions. (H) Example song spectrograms (0.5 s) extracted at moments when neural sequences were detected in an isolated bird (older juvenile, 64 dph) (I) Correlation of these sequences with eight song features (top to bottom: amplitude, entropy, pitch goodness, aperiodicity, mean frequency, pitch, frequency modulation, amplitude modulation). Factor duration 0.5 s, indicated by colored bars above, triangle at center. (J) Strength of song locking (see Methods). (K) Percent of the song covered by some sequence. (L) Example of sequence abnormalities in an isolated bird (same as in E). Sequences of inconsistent length (8/8 isolated birds) and ensemble persistent activity (7/8 isolated birds) are annotated in red.

By observing the neural activity in HVC of isolated birds, we found that the HVC network activity can mature into long repeatable sequences even without exposure to a tutor. However, there are some key differences between typical adult HVC sequences and those found in isolated birds, suggesting which features of HVC development rely on exposure to a tutor. Next, we observe HVC in isolated birds immediately before and after delayed exposure to a tutor. Birds isolated from a tutor are able to learn a song if exposed to a tutor before the end of a critical period, typically around age 65 days post hatch (dph), but are increasingly unable to learn at later ages [24, 25, 26]. Although only half of our late tutored birds successfully learned from the tutor, we observed an interesting correlation between HVC activity prior to tutoring and the degree to which birds learned. Namely, birds with highly song-locked HVC activity prior to tutoring typically failed to learn, while birds with less song-locked activity tended to learn. In the birds that did learn, we were able to track sequences throughout the course of learning. Pre-existing self-organized HVC sequences persisted throughout major changes to the song, forming a substrate for newly learned song elements. Together, these results point at how the brain may self-organize, and at the interplay between self-organized structure and the ability to incorporate new information from a tutor.

## 2 Results

### 2.1 Neural sequences are present in isolated birds, but atypical

We first asked whether the songs of isolated birds involve the same neural path-ways and neuronal sequences responsible for generating typical song. We carried out functional calcium imaging of large populations of neurons in HVC of isolated birds at a range of ages. Sequences of neuronal activity in HVC have previously been analyzed by aligning neuronal activity to repeatable elements of the song [27], an approach with limited utility in isolated birds due to the high variability of their songs. Instead, we extract neural sequences directly from the calcium signals using an unsupervised algorithm [28] to find the sequences that best fit the neural data. This technique reveals the existence of significant sequential activity in HVC of isolated birds (Figure 1B,C). It also reveals long continuous sequences in data acquired from typical adult HVC (Figure 1D) as expected from previous work [27, 19, 18].

The sequences found in isolated birds are surprisingly typical in some respects, but atypical in others, especially in their correlation to vocal output. As in typical HVC sequences, neurons in isolated birds participate at characteristic moments during the sequence (Figure 1E), and many neurons participate in at least one sequence (Figure 1F). Neurons that participate in a sequence tend to fire at a majority of sequence occurrences (Figure 1G). Neural sequences are correlated with precisely timed song features in isolated birds’ song (Figure 1H, I, song features calculated as in [29]). However, song locking in isolated birds was only on average 0.58 times as strong as in a typically tutored adult bird (Figure 1J, see Methods). Finally, in isolated birds, on average only 61% of each song bout is represented by a detected HVC sequence, substantially less than the complete sequence coverage found in typically tutored birds [19, 18, 17] (Figure 1K, see Methods). HVC activity in isolated birds exhibits additional qualitative differences from that in typically tutored birds. While HVC neurons generate only brief bursts of spikes in tutored birds, neurons in isolated birds sometimes generated extended periods of continuous activity, especially during long syllables of variable duration (Figure 1L, 7/8 birds exhibited multiple instances of persistent activity, coordinated across at least 3 neurons, and lasting at least 500ms). This contrasts with long syllables of typical adult song which are all generated by extended sequences of brief bursts. In addition, HVC sequences in isolated birds exhibit variable durations, often truncating at different points (Figure 1L, 8/8 birds), producing syllables of highly variable duration. Such truncations in the middle of a syllable sequence are very unusual in typically tutored birds [30]. These atypical modes of HVC activity suggest several possible mechanisms to understand characteristic features of isolate song, abnormally long syllables and those of variable duration [12]. For example, syllables in isolated birds may exhibit variable duration when their underlying HVC sequences are truncated at different points.

We wondered if the existence of sequences in HVC of socially isolated birds occurs only after the closure of the critical period (i.e. a product of an already atypical isolate song) or whether they develop at an even earlier age when birds have not yet heard a tutor song, but can still be tutored. We recorded in 5 birds at ages 57-64 dph, prior to tutor exposure, and found strong evidence for HVC sequences (Figure S1A). There was not a significant correlation between the age of the bird and any sequence features we measured (Figure S1B-F, linear regression model, significance threshold p < 0.5, comparing to a constant model). The correlations were not significant both when we restricted to birds within the traditional critical period (<65 dph), and when we included data from three older isolated birds (68-117 dph). Thus, the large (several fold) bird-to-bird variability in sequence properties (Figure S1B-F) is not explained by age, and likely due to inter-individual variability in developmental timecourses.

### 2.2 Prior to tutoring, birds that will learn exhibit HVC sequences that are relatively immature and decoupled from vocal output

Next, we asked whether properties of the HVC sequences relate to the ability of birds to learn a new song from a tutor. Many of our young isolated birds were eventually tutored at an age around the critical period and we found that half of them learned elements of their tutor song, while the others developed fully isolate song. We classified birds as learners if their song had an Imitation Score metric [31] greater than 0.5. The songs of non-learners remained highly variable and isolate-like even after tutoring (Figure 3A). In contrast, learner birds developed a new syllable within a day or two after tutoring, and ultimately sang typical adult song, consisting of stereotyped motifs (Figure 3B).

An analysis of HVC activity revealed that sequences prior to tutoring were systematically less mature/‘crystallized’ in birds that learned than in birds that failed to learn. Learner birds had fewer sequences than non-learners (Figure 2C, average 2 sequences in learners, 3.25 sequences in non-learners, p = 0.029, Wilcoxon rank sum test). Sequences in learner birds were more weakly correlated to song features (Figure 2D, average 0.20 s learner, 0.55 s non-learner, p = 0.0034, Wilcoxon rank sum test). Sequences in learners had lower autocorrelation, a measure of how repeatably/rhythmically they are produced [17], than non-learners (Figure 2E, average 0.125 s learner, 0.244 s non-learner, p = 0.018, Wilcoxon rank sum test). Three additional measures of sequence maturity, all related to intrinsic sequence properties were calculated. While non-learners also trended higher in these measures, the differences were not significant (Wilcoxon rank sum tests, Neural participation: average 45% learner, 70% non-learner, p = 0.2; Reliability: average 69% learner, 74% non-learner, p = 1; Coverage: average 51% learner, 71% non-learner, p = 0.34).

**Figure 2:**
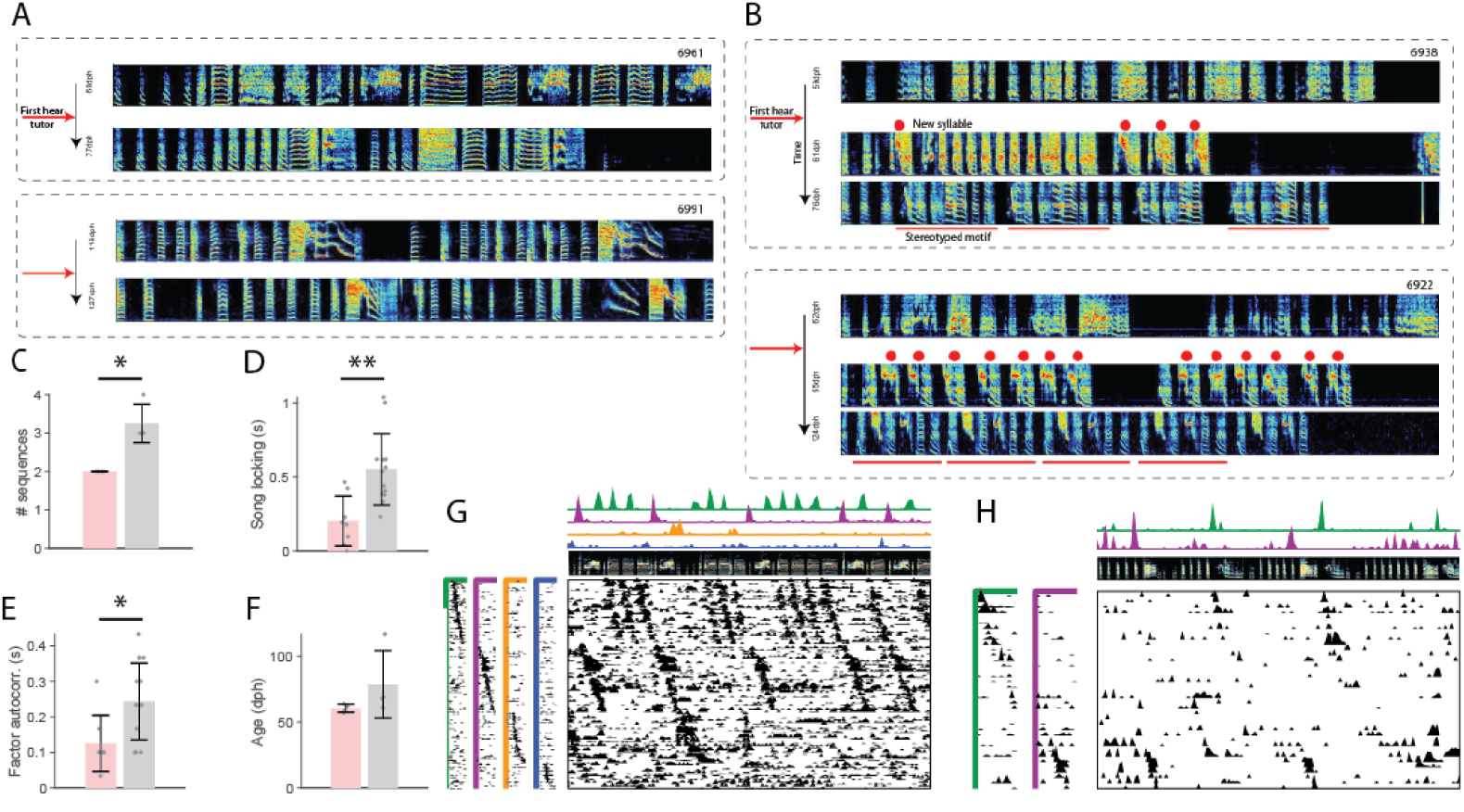
Relation between HVC sequence maturity and subsequent song learning (A) Example spectrograms for two non-learner birds, prior to tutoring and several weeks later (at least 77dph). (B) Example spectrograms for two learners, prior to tutoring, shortly after tutoring, and several weeks later. Red dots mark the new syllable. Red bars mark stereotyped motif. (C-E) Three measures of HVC sequence maturity for learners (pink) and non-learners (gray). Error bars denote standard deviation (* : p<0.05, ** : p<0.01). (c) Number of sequences in HVC. (D) Fraction of neurons that participate in a sequence. (E) Autocorrelation of sequence factor timecourses. (F) Age of first tutoring for learners and non-learners. (G-H) Example pre-tutoring data from two birds that were brothers. (G) A non-learner, first tutored at 61 dph. (H) A learner, first tutored at 64 dph.

The age of tutoring was not significantly correlated with whether the bird was a learner or non-learner (Figure 3F, average 60.5 dph learner, 78.75 dph non-learner, p = 0.11, Wilcoxon rank sum test). For example, one of the younger birds in our dataset (61 dph) was a non-learner, and had particularly clear HVC sequences before tutoring (Figure 3G). This bird’s brother, tutored 3 days later, was a learner, and had sequences that appear far less mature (Figure 3H). Together, these results suggest that the presence, at the time of tutoring, of robust song-locked sequences, may inhibit learning. In other words, learning may be better supported by more immature sequences that are more independent from vocal output.

**Figure 3:**
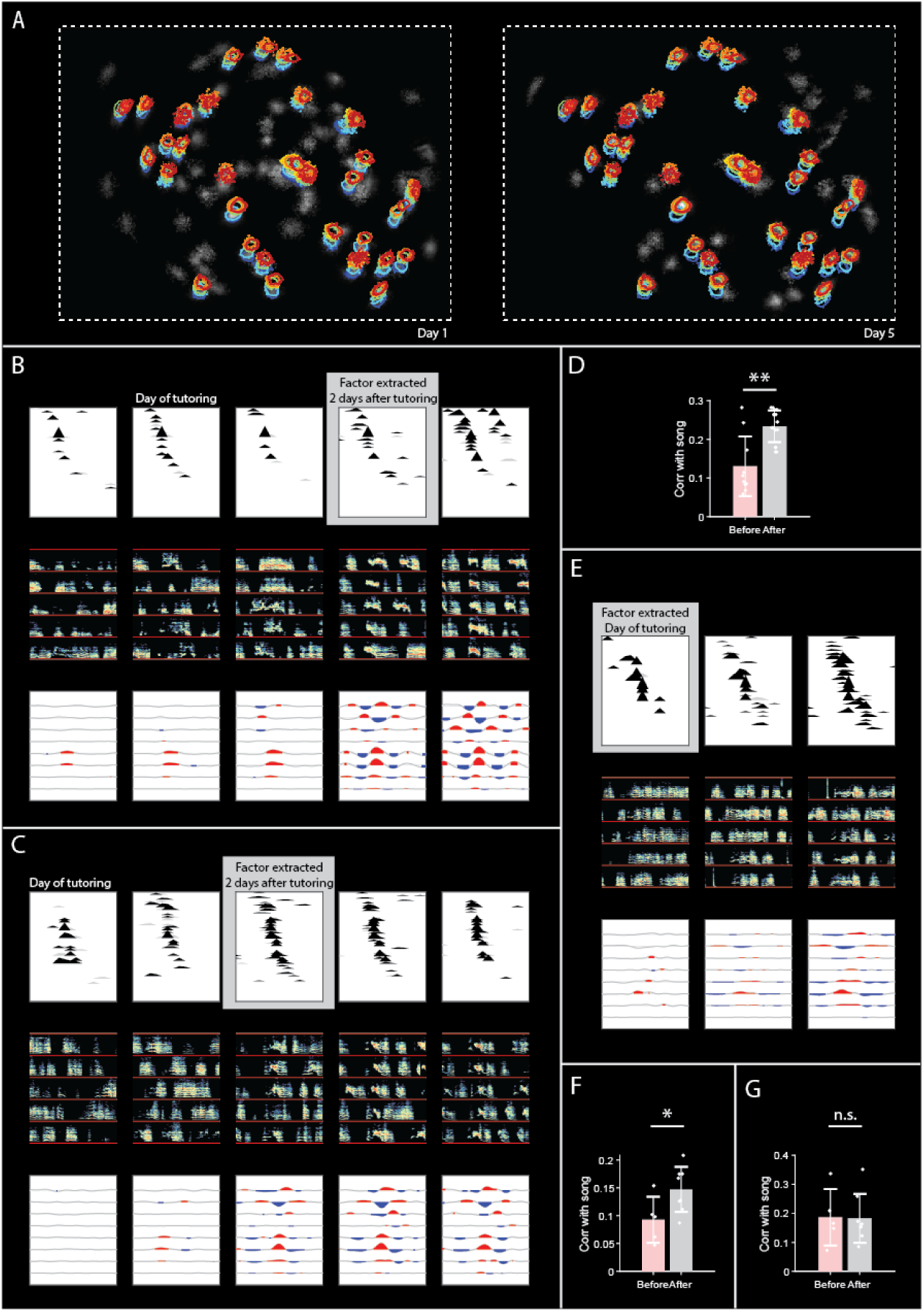
Tracking HVC sequences as isolated birds rapidly learn a new syllable (A) Neurons detected before (left) and after (right) tutoring shown in grayscale (CNMF E algorithm). Colored contours indicate locations of neurons tracked across five days, from blue to red (B) Sequence in HVC, tracked before and after first tutor exposure (see Methods), through the development of a new syllable. Sequence extracted from data two days after tutoring, and neurons sorted according to participation in this sequence. (Top) On each recording day, cross correlation of neurons with the sequence that becomes associated with the new syllable. Significant bins are shown in black, non-significant bins in gray (p=0.05, Bonferroni corrected, compared to circularly-shifted control) (Middle) On each recording day, example spectrograms at times when the sequence occurs on each day (Bottom) On each recording day, cross-correlation of sequence with acoustic features (amplitude, entropy, pitch goodness, aperiodicity, mean frequency, pitch, frequency modulation, and amplitude modulation) (C) Same as B for a different example bird. (D) Correlation with song amplitude before (pink) and after (gray) tutoring for all sequences in learner birds extracted data when a new syllable had been learned. (E) Similar to B and C, for a different example bird. Here sequences are extracted from pre-tutoring data, then tracked forward in time. (F) Song locking (maximum cross-correlation with song amplitude) before and after tutoring for the pre-tutoring sequences that had weaker song locking. (G) Song locking before and after tutoring for the pre-tutoring sequences that started off with stronger song locking.

### 2.3 Tracking HVC sequences across rapidly learned song changes

In late-tutored birds that learned, the speed with which new syllables appeared was striking. These birds developed a new syllable within a day or two after tutoring (Figure 2A, B), as has been previously described [32, 33, 34]. These new syllables appeared to emerge *de-novo*, not by syllable differentiation as is common in tutored birds.

We wondered if these birds, which learned a new syllable rapidly after tutoring, formed a *de-novo* HVC sequence for this new syllable, or perhaps used a pre-existing sequence. We were able to track neurons in our calcium imaging data throughout the course of tutoring (Figure 3A, see Methods, Gu et al., in preparation), enabling us to see what happens to neural activity during rapid changes in the song. We first extracted neural sequences associated with new post-tutoring syllables, then followed these neurons back in time to find that the sequence existed even prior to tutoring (Figure 3B,C, see Methods). However, the sequence prior to tutoring was surprisingly ‘latent’. That is, the sequence was relatively uncoupled to vocal output, without a strong correlation to song syllables. Combining data from the four birds that learned a new syllable rapidly after tutoring, neural sequences extracted two days after tutoring appeared to become more song locked after tutoring (Figure 3D, p=0.0048, Wilcoxon rank sum test).

Next, we aimed to control for the possibility that the appearance of sequences becoming progressively more locked to vocal output after tutoring was due to the fact that sequences were extracted from neural data recorded after tutoring. We directly extracted HVC sequences from exclusively pre-tutoring neuronal data and tracked them forward in time until a new syllable appeared. Sequences that were initially relatively ‘latent’ persisted, becoming progressively more correlated with vocal output, ultimately tightly locked to a new syllable (Figure 3E). Each of the ‘learner’ birds appeared to have two HVC sequences present prior to tutoring. Of these sequences, the ones that started off less correlated with song amplitude exhibited a significant increase in correlation with song amplitude after tutoring (Figure 3F, p=.045, Wilcoxon rank sum test). The sequences that started off more correlated with song amplitude did not significantly change their correlation with song amplitude (Figure 3G, p=1, Wilcoxon rank sum test). Together, these results are consistent with the view that the emergence of new syllables after tutoring may co-opt existing HVC sequences, including relatively ‘latent’ sequences.

## 3 Discussion

We set out to determine whether the formation of sequences in HVC depends on prior exposure to a tutor song. By observing the neural activity in HVC of isolated birds, we found that HVC network activity can form long repeatable sequences even in birds that had no prior exposure to vocal tutoring. Sequences in isolate HVC exhibit some properties of typical HVC, with many neurons reliably participating in sequences, and sequences being correlated to vocal output. However, sequences in isolated birds were less reliable and less tightly correlated with vocal output than has been described in typical birds, and exhibited abnormal truncations and persistent activity.

We had previously hypothesized that the experience of hearing a tutor may seed the formation of HVC sequences of the appropriate number and durations [35], but our new data reveal that HVC sequences exist even prior to tutoring. Thus, there must be a way for sequences to form without the prior storage of a tutor memory. In models of Hebbian learning in HVC, sequences can form in networks driven by random inputs rather than patterned inputs [22]. However, in this case the distribution of sequence durations no longer matches syllable durations found in typical adult birds, but is instead more consistent with the highly variable and atypically long syllables that occur in birds that have never heard a tutor (isolate song) [12, 10]. Thus our findings may be consistent with the view that sequences can emerge in isolate birds by a combination of simple Hebbian learning mechanisms together with spontaneous activity either within HVC or driven by the inputs to HVC.

Our discovery of latent sequences suggests a separation between neural processes for building a stable representation of states within a task (i.e., sequential moments in time), and neural processes for associating an action with each state. Thus, sequences may gradually emerge in the maturing HVC network via simple Hebbian processes [23, 22, 17], but may remain relatively decoupled from downstream motor neurons until a memory of the tutor song is learned and reinforcement learning processes begin.

From a computational perspective, what do latent sequences tell us about how the brain learns? By latent sequences, we mean sequences that are initially only weakly correlated with vocal output, but are subsequently used to produce learned song changes. In reinforcement learning models of song learning, HVC sequences remain relatively stable even as the song changes [36, 37, 38, 39], consistent with our observation of stable sequences. This is in contrast with other models of song learning, like the ‘inverse model’ [40, 41, 42]. In the inverse model, each motor neuron produces the same vocal output at different times during vocal learning; song changes are caused by pre-motor neurons (e.g. HVC) being activated in a different order. In contrast, we observed relatively stable sequences throughout learned song changes. Our results are consistent with data from primary motor cortex of macaques operating a brain-computer interface—a fixed repertoire of activity patterns are associated with different movements after learning [43]. Our results are also consistent with the idea that the brain may use pre-existing sequential patterns to rapidly learn from new experience, for example the existence of sequences in the hippocampus prior to exposure to new environments [44, 45, 46, 47, 48].

If the brain is able to build on latent structure to learn from sparse data, essentially implementing inductive bias, we might expect different forms of latent structure for different tasks. Zebra finches are known to develop typical songs, including typical syllable durations, after being tutored by atypical isolate songs, relying on species-specific ‘priors’ to achieve species-typical syllable durations. The latent sequences we observed tended to last on the order of a hundred milliseconds—the same as the duration of typical zebra finch syllables. Might other species that sing faster songs (e.g. grasshopper sparrow) or slower songs (e.g. white-throated sparrow) exhibit latent sequences of shorter or longer durations? One might imagine that the speed of latent sequences could be genetically specified by expression levels of ion channels with different time constants within HVC. Alternatively, the duration of latent sequences could be specified by the amount of time it takes for HVC to get feedback from respiratory and/or auditory centers, which may also have their own intrinsic rhythmicity [49, 50, 51]. Each of these possible sources of latent HVC structure could be tested in further experiments. By whatever mechanism latent sequences arise, they appear to be capable of supporting song learning, at least in the case of delayed tutoring. More generally, the ability of brains to generate complex learned behavior may depend on the intrinsic developmental formation of appropriate latent dynamics in motor and sensory circuits.

## 4 Materials and Methods

### 4.1 Table of key resources for imaging HVC sequences

Key resources, and references for how to access them, are listed in Table 1.

**Table 1:** Links to key resources used for measuring HVC sequences during rapid learning

| Software/algorithm | Source | Link to code |
| --- | --- | --- |
| seqNMF | [28] | https://github.com/FeeLab/seqNMF |
| CNMF_E (cell extraction) | [52] | https://github.com/zhoupc/CNMF_E |
| STAT (tracking neurons across days) | Gu et al., in preparation | will post preprint and submit |
| Chronux (spectrogram computation) | [53] | http://chronux.org/ |
| SAP (Sound Analysis Pro) | [29] | http://soundanalysispro.com/ |
| SI (Song Imitation) | [31] | https://doi.org/10.1371/journal.pone.0096484 |
| MATLAB | MathWorks | www.mathworks.com |
| Dataset | Source | Link to data |
| HVC, rapid learning | This paper | will post |
| Other | Source | Link |
| Zebra finches (*Taeniopygia guttata*) | MIT animal facility |  |
| AAV9.CAG.GCaMP6f.WPRE.SV40 | [54] | https://pennvectorcore.med.upenn.edu |
| Miniature microscope | Inscopix nVista | https://www.inscopix.com/nvista |

### 4.2 Animal care and use

For this study, Imaging data was collected in 9 male zebra finches (*Taeniopygia guttata*) from the MIT zebra finch breeding facility (Cambridge, MA). Animal care and experiments were carried out in accordance with NIH guidelines, and reviewed and approved by the Massachusetts Institute of Technology Committee on Animal Care.

In order to control exposure to a tutor song, 8 birds were foster-raised by female birds, which do not sing, starting on or before post-hatch day 15 (15 dph). Starting between 40 dph and 50 dph, these birds were housed singly in custom-made sound isolation chambers. An additional bird was tutored by his father, as is typical. After a couple of days of acclimation to the lab environment, birds were anesthetized with isoflurane, and were given a surgery to inject virus to express the functional indicator GCaMP6f and implant a GRIN (gradient index) lens (see below). Analgesic (Buprinex) was administered 30 min prior to the surgery, and for 3 days postoperatively. After at least a week for virus expression, an Inscopix miniscope baseplate was attached to the existing implant. Birds were acclimated to the miniscope for several days. Once birds started singing with the miniscope, functional calcium signals were recorded for several days. To avoid photobleaching, short files (approximately 10 seconds) were obtained, typically fewer than 50 files per day. Once some pre-tutoring singing data had been obtained, birds were tutored briefly (5-10 song bouts from a tutor bird) each day.

### 4.3 Expression of functional calcium indicator GCaMP6f

The calcium indicator GCaMP6f was expressed in HVC by intercranial injection of the viral vector AAV9.CAG.GCaMP6f.WPRE.SV40 [54] into HVC. In the same surgery, a cranial window was made using a relay GRIN (gradient index) lens (1mm diamenter, 4mm length, Inscopix) implanted on the surface of the brain, after the dura was removed. After at least one week, in order to allow for sufficient viral expression, recordings were made using the Inscopix nVista miniature fluorescent microscope.

### 4.4 Extraction of neuronal activity and background subtraction using CNMF E

Neuronal activity traces were extracted from raw fluorescence movies using a constrained non-negative matrix factorization algorithm, CNMF_E, that is specialized for microendoscope data by including a local background model to remove activity from out-of-focus cells [52]. Custom software (Shijie Gu, Emily Mackevicius, Pengcheng Zhou) was used extend the CNMF E algorithm to combine batches of short files (BatchVer) and track individual neurons over the course of multiple days (STAT, Gu, et. al., in preparation, see below).

### 4.5 Unsupervised discovery of neural sequences using seqNMF

We addressed the challenge of needing to detect neural sequences in HVC without relying on aligning neural activity to the song by developing an unsupervised algorithm, seqNMF [28]. This was necessary because juvenile songs are highly variable and difficult to parse into repeatable syllables, and because we wanted to allow for the possibility that HVC activity might be more stereotyped than the song. Briefly, seqNMF factorizes data into exemplar sequence factors (**W**’s). Each sequence factor has a corresponding timecourse (**H**). Convolving each exemplar with its respective timecourse produces an approximate reconstruction of the original data 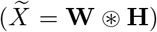. SeqNMF returns a factorization that minimizes reconstruction error, subject to a penalty term that encourages simpler factorizations.

### 4.6 Preprocessing calcium traces prior to running seqNMF

We performed several preprocessing steps before applying seqNMF to functional calcium traces extracted by CNMF E. First, we estimated burst times from the raw traces by deconvolving the traces using an AR-2 process. The deconvolution parameters (time constants and noise floor) were estimated for each neuron using the CNMF E code package [52]. Some neurons exhibited larger peaks than others, likely due to different expression levels of the calcium indicator. Since seqNMF would prioritize the neurons with the most power, we renormalized by dividing the signal from each neuron by the sum of the maximum value of that row and the 95*th* percentile of the signal across all neurons. In this way, neurons with larger peaks were given some priority, but not much more than that of neurons with weaker signals.

### 4.7 Estimating the number of significant sequences in each dataset

The number of sequences present in real neuronal datasets can be slightly ambiguous, so we used several methods to arrive at and validate an estimate for the number of significant neural sequences present in each dataset. It is important to note that, since our datasets are short, there may be additional neural sequences in HVC that do not appear, or do not achieve significance, in our datasets. In order to cross-validate sequences on held-out data, we split each dataset into a training set (75%) and a test set (25%). Sequences were detected in the training set, and significance was measured in the test set by assessing how much the overlap of the sequences with the test data compared to null (time-shifted) sequences. In order to choose a value for the seqNMF parameter *λ* that balances reconstruction cost with correlation (redundancy) cost, we swept *λ* with *K* = 10 and *L* = 0.5 seconds to find *λ*0, the cross-over point that balances these cost terms (Figure S2A). Based on analysis on simulated data [28], where values of *λ* at or slightly above *λ*0 yielded the correct number of sequences, we looked at the distribution of significant sequences at *λ* = *λ*0 and *λ* = 2*λ*0 (Figure S2B), and chose as our estimate a number between the peaks of these two distributions. We validated these estimates in two ways. First, we ran seqNMF with K equal to this estimate and *λ* = 0, and confirmed that the resulting sequences tended to be significant on held-out data. Next, we ran seqNMF on the entire dataset at this K from 25 different random initial conditions, and confirmed that the sequences were consistent across the different runs (Figure S2C). Consistency measures the extent to which there is a one-to-one mapping between the factors of two different factorizations [28]. When this analysis was run at a K higher than the estimated K, results tended to be less consistent (Figure S2D).

### 4.8 Selecting a consistent factorization

For each dataset, we selected the most consistent factorization on which to perform all further analysis. Once we had selected an appropriate number of sequences for each dataset, using the analyses described above, we ran seqNMF 25 times at this value of *K* from different random initial conditions, and picked the factorization that was most consistent with the other factorizations (Figure S2D). Factorizations at *K* chosen by the above methods tended to be more consistent than factorizations at higher *K* (Figure S2D).

### 4.9 Significance testing for cross-correlation analyses

Several of our results involve analyzing the temporal relationship between different timecourses (factors and neurons; factors and song acoustic features; factor autocorrelations). These analyses involve testing the significance of the cross-correlation between two timeseries, compared to null cross-correlation values that could occur if the signals were circularly shifted relative to each other by a random large timelag. Before measuring cross-correlations, we centered each signal to have zero mean. If we are assessing the cross-correlation at lags in the range from -L to L, we want to compare values measured here to null values measured at random lags longer than L. We compute the cross-correlation at each lag *ℓ* in the range −*T* < *ℓ* < *T*, where T is the length of the timeseries, by circularly shifting one of the timeseries by *ℓ* and computing the dot product with the other timeseries. We then use the cross-correlations at null lags (−(*T*− *L*) < *ℓ* < −*L* or *L* < *ℓ* < (*T*− *L*)) to determine a Bonferroni-corrected significance threshold. The threshold is the 100 × (1− *p/Num*)^*th*^ percentile of the absolute value of these null cross-correlations, where *Num* is the number of comparisons (2L times the number of tests being run), and *p* is the *p*-value. Significance is achieved for lags at which the measured cross-correlation exceeds this value.

### 4.10 Assessing song locking, the cross-correlation between each factor and acoustic song features

Several of our results involve quantifying the temporal relationship between sequence timecourses (**H**’s) and the song. To do this, we measured the cross-correlation of sequences with song acoustics using 8 acoustic features common in the songbird literature [29]: amplitude, entropy, pitch goodness, aperiodicity, mean frequency, pitch, frequency modulation, and amplitude modulation. Each of these acoustic features is measured from the song at 1ms resolution using standard software (Sound Analysis Pro, http://soundanalysispro.com/, [29]). The seqNMF H’s are upsampled to this resolution, then cross-correlation between each **H** and each song feature is assessed using the above procedure, with L = 1 second, p = 0.05, and Bonferroni correction (2000 timebins) x (8 features) x (K sequences). The overall measure of song locking is computed by integrating the number of seconds that a given sequence has significant correlation with each of the song features.

### 4.11 Assessing which neurons participate in each sequence

Several of our results involve assessing which neurons participate in each sequence. In order to do this, we measure whether there is a significant cross-correlation between each neuron and each factor (with L=0.5 seconds, p = 0.05, and Bonferroni correction (30 timebins) x (N neurons) x (K sequences)). Note that, since seqNMF is run on the neural data, it is guaranteed that some neurons will be correlated with the factors —the primary aim of this test is to assess which neurons are in which sequences.

### 4.12 Tracking HVC projection neurons over the course of major song changes

A core motivation for using calcium imaging methods instead of other methods was the possibility to track HVC projection neurons over the course of major song changes. HVC projection neurons are particularly difficult to record with electrophysiological methods—current methods are unable to record an HVC projection neuron for more than a few hours, and tend to record one, or at most three, projection neurons at a time [55, 17]. Previous studies of song-locked HVC activity throughout the learning process could only track changes in the neural population that occurred at a timescale slower than a week, because population statistics had to be compiled from single-neuron recordings [17]. This technique misses rapid changes that can happen within a day [32], and is unable to assess the stability of HVC sequences.

Stability of HVC sequences over time can be assessed using calcium imaging, though some challenges remain due to the potential for errors in tracking neurons across days. Single-photon calcium imaging methods have been used to address the stability of HVC sequences in adult birds with stable songs, observing stable song-locked activity in slightly more than half of HVC projection neurons, and unstable song-locked activity in slightly less than half of HVC projection neurons [56]. This measure is likely an underestimate of the stability of HVC activity, since noise in tracking cell locations across days could lead to perceived instability. Thus, HVC sequences appear relatively stable in birds with stable song, but what about birds whose songs are changing? The potential for errors in tracking neurons across days was one factor in our decision to record in birds undergoing very rapid learning. It was necessary for us to expand upon previous methods for tracking neurons recorded by calcium imaging over time [57], likely due to the relatively short individual file sizes in our dataset from singing juvenile birds (we recorded many short files each day, when the birds happened to sing, instead of longer continuous files).

We tracked the activity of populations of HVC neurons over multiple days using Spatial Tracking Across Time (STAT, Gu et al., in preparation). This method builds off of previous methods [57], where individual cell pairs’ shape spatial correlation and distance are used to determine the correspondences between cells extracted from different sessions. STAT also considers local neighborhood motion consistency in computing the optimal tracking of cells across sessions, and requires less manual supervision. The local motion consistency is optimized using the Hungarian Method, a combinatorial optimization algorithm that solves assignment problems in polynomial time. Cells that have no good match are excluded, as are cells with abnormal coefficient of variations. Finally, the results of the matching algorithm are checked manually.

### 4.13 Tracking sequences extracted on one subset of a dataset to another subset of the dataset

In order to track a sequence, **W**, extracted in one subset of a dataset (**X**_1_, for example before tutoring) to another subset of the dataset (**X**_2_, for example after tutoring), we first mean-subtract **W** and **X**_2_ along the time dimension, then estimate 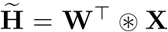. In order to assess whether a neuron significantly participates in **W** in dataset **X**_2_, we bootstrap using control datasets 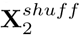, in which data from each neuron is circularly shifted in time by a different random amount. We then ask whether the neuron participates more strongly in the real dataset compared to participation calculated on control datasets (p=0.05 significance threshold, Bonferroni corrected for the number of neurons and the number of time-lags). Specifically, we compare 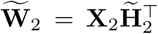 to 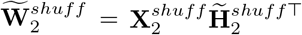.

## 5 Assessing sequence coverage of song bouts

Sequence coverage quantifies the observation that sequences in isolated birds appear to pop on and off at somewhat arbitrary moments in bouts, leaving some sections of some bouts with no clear sequences present. First, the moments when each sequence occurs is estimated by computing when 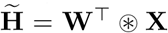 is larger than expected by chance (Bonferroni-corrected 95% percentile of 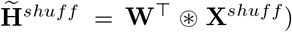. Next, the sequence is convolved with the corresponding **W**. Finally, the total number of seconds when some sequences was present is divided by the total number of seconds in the bout, and multiplied by 100, to get the percent of the bout covered by some sequence. Note that sequence coverage is distinct from previously described measures of burst coverage within a repeatable adult song motif [18].

## 6 Acknowledgements

This work was supported by a grant from the Simons Collaboration for the Global Brain, the National Institutes of Health (NIH) [R01 DC009183] and the G Harold and Leila Y Mathers Charitable Foundation. ELM received support through the NDSEG Fellowship program and the Simons Society of Fellows. Special thanks to Andrew Bahle for comments on earlier versions of the manuscript.

## Supplementary figures

**Figure S1:**
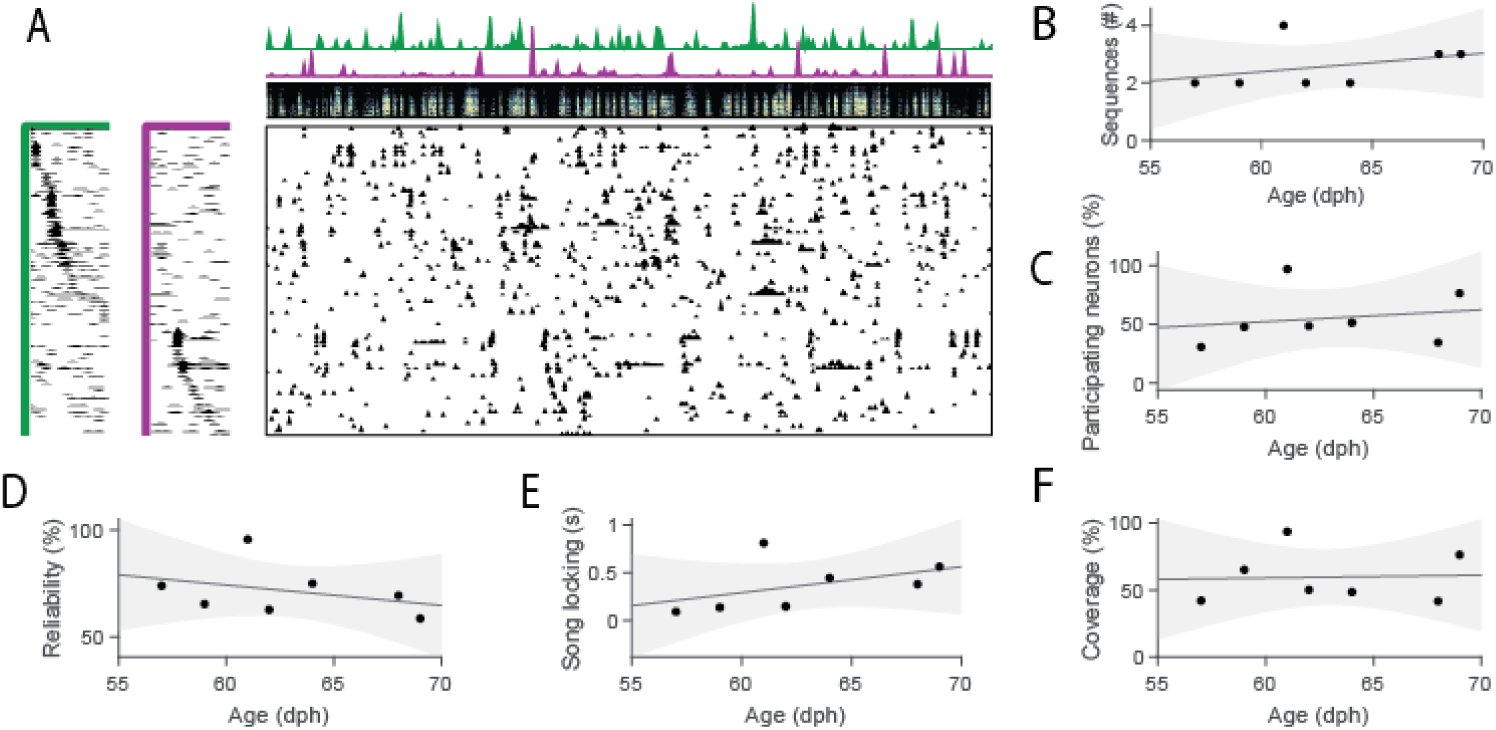
HVC sequences exist even in young isolated birds (A) Example HVC sequences recorded in a young isolated bird (59 dph) (B-F) Sequence properties as a function of age in 7 juvenile isolated birds (5 birds recorded prior to the closing of the traditional critical period (<65 dph), and 2 older juvenile birds (65 dph - 90 dph)). Line denotes least squares fit, gray area 95% confidence interval. (B) Number of HVC sequences extracted. (C) Percent of neurons participating in at least one sequence. (D) Reliability of neural participation across sequence renditions. (E) Song locking. (F) Percent of the song covered by at least one sequence.

**Figure S2: Supplementary Figure 1.**
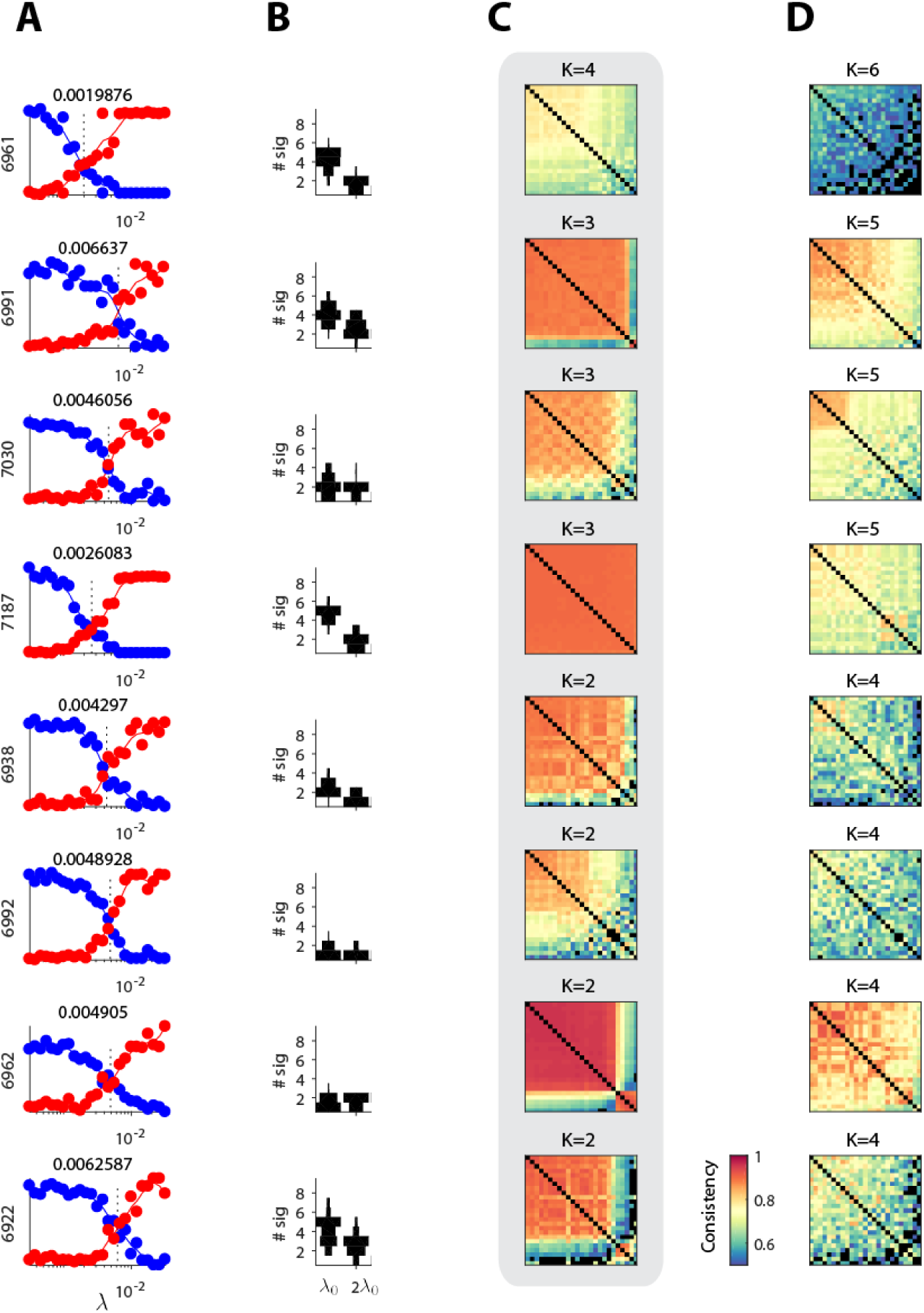
Estimating the number of significant sequences in each dataset (A) Reconstruction cost (red) and correlation cost (blue) as a function of *λ* (with K=10, L=0.5 seconds) for 8 datasets (pre-tutoring data from 8 different birds). The crossover point, *λ*_0_, is stated and marked by a dashed line. (B) Histogram of the number of significant sequences at *λ*_0_ and 2*λ*_0_ for these datasets. (C) For the chosen K, and *λ* = 0, consistency across 25 runs of seqNMF from different random initializations. Factorizations are sorted from most to least consistent. (D) Consistency matrix for 25 runs at K above the estimated K.

## Notes

### Competing Interest Statement

The authors have declared no competing interest.

## References

[1] Noam Chomsky. On nature and language. Cambridge University Press, 2002.

[2] Torsten N. Wiesel and David H. Hubel. Single-cell responses in striate cortex of kittens deprived of vision in one eye. Journal of Neurophysiology, 26(6):1003–1017, 1963. PMID: 14084161.

[3] M. Bear and W. Singer. Modulation of visual cortical plasticity by acetylcholine and noradrenaline. Nature, 320:172–176, 1986.

[4] Brandon J. Farley, Hongbo Yu, Dezhe Z. Jin, and Mriganka Sur. Alteration of visual input results in a coordinated reorganization of multiple visual cortex maps. Journal of Neuroscience, 27(38):10299–10310, 2007.

[5] Jie Ye, Priti Gupta, Pragya Shah, Kashish Tiwari, Tapan Gandhi, Suma Ganesh, Flip Phillips, Dennis Levi, Frank Thorn, Sidney Diamond, Peter Bex, and Pawan Sinha. Resilience of temporal processing to early and extended visual deprivation. Vision Research, 186:80–86, 2021.

[6] Kathryn L Hildyard and David A Wolfe. Child neglect: developmental issues and outcomes. Child Abuse and Neglect, 26(6):679–695, 2002.

[7] I. Moreno-Torres, S. Madrid-Cánovas, and G. Blanco-Montañez. Sensitive periods and language in cochlear implant users. Journal of child language, 43:479–504, 2016.

[8] A. Kral, M. F. Dorman, and B. S. Wilson. Neuronal development of hearing and language: Cochlear implants and critical periods. Annual review of neuroscience, 42:47–65, 2019.

[9] Ofer Tchernichovski and Gary Marcus. Vocal learning beyond imitation: mechanisms of adaptive vocal development in songbirds and human infants. Current Opinion in Neurobiology, 28:42–47, 2014.

[10] Philip H Price. Developmental Determinants of Structure in Zebra Finch Song. 93(2):260–277, 1979.

[11] Heather Williams, Kerry Kilander, and Mary Lou Sotanski. Untutored song, reproductive success and song learning. Animal Behaviour, 45(4):695–705, apr 1993.

[12] Olga Fehér, Haibin Wang, Sigal Saar, Partha P Mitra, and Ofer Tchernichovski. De novo establishment of wild-type song culture in the zebra finch. Nature, 459(7246):564–568, may 2009.

[13] Mugdha Deshpande, Fakhriddin Pirlepesov, and Thierry Lints. Rapid encoding of an internal model for imitative learning. Proceedings of the Royal Society of London B: Biological Sciences, 281(1781), 2014.

[14] Lucy A Eales. Do zebra finch males that have been raised by another species still tend to select a conspecific song tutor? Animal Behaviour, 35(5):1347–1355, 1987.

[15] Mets D.G. and Brainard M.S. Genetic variation interacts with experience to determine interindividual differences in learned song. Proc Natl Acad Sci, 115(2):421–426, January 2018.

[16] David G Mets and Michael S Brainard. Learning is enhanced by tailoring instruction to individual genetic differences. Elife, 8:e47216, 2019.

[17] Tatsuo S. Okubo, Emily L. Mackevicius, Hannah L. Payne, Galen F. Lynch, and Michale S. Fee. Growth and splitting of neural sequences in songbird vocal development. Nature, 528(7582):352–357, nov 2015.

[18] Galen F Lynch, Tatsuo S Okubo, Alexander Hanuschkin, Richard HR Hahnloser, and Michale S Fee. Rhythmic continuous-time coding in the songbird analog of vocal motor cortex. Neuron, 90(4):877–892, 2016.

[19] Michel A. Picardo, Josh Merel, Kalman A. Katlowitz, Daniela Vallentin, Daniel E. Okobi, Sam E. Benezra, Rachel C. Clary, Eftychios A. Pnevmatikakis, Liam Paninski, and Michael A. Long. Population-Level Representation of a Temporal Sequence Underlying Song Production in the Zebra Finch. Neuron, 90(4):866–876, may 2016.

[20] G. Kosche, D. Vallentin, and M. A. Long. Interplay of Inhibition and Excitation Shapes a Premotor Neural Sequence. Journal of Neuroscience, 35(3):1217–1227, jan 2015.

[21] D. Vallentin, G. Kosche, D. Lipkind, and M. A. Long. Inhibition protects acquired song segments during vocal learning in zebra finches. Science, 351(6270):267–271, jan 2016.

[22] Ila R Fiete, Walter Senn, Claude Z H Wang, and Richard H R Hahnloser. Spike-time-dependent plasticity and heterosynaptic competition organize networks to produce long scale-free sequences of neural activity. Neuron, 65(4):563–76, feb 2010.

[23] Joseph K Jun and Dezhe Z Jin. Development of neural circuitry for precise temporal sequences through spontaneous activity, axon remodeling, and synaptic plasticity. PloS one, 2(8):e723, jan 2007.

[24] Sarah E. London. Developmental song learning as a model to understand neural mechanisms that limit and promote the ability to learn. Behavioural Processes, 163:13–23, jun 2019.

[25] Sharon M.H. Gobes, Rebecca B. Jennings, and Rie K. Maeda. The sensitive period for auditory-vocal learning in the zebra finch: Consequences of limited-model availability and multiple-tutor paradigms on song imitation. Behavioural Processes, 163:5–12, jun 2019.

[26] Klaus Immelmann. Song development in the zebra finch and other estrildid finches. Bird vocalizations, pages 61–77, 1969.

[27] Michael A Long, Dezhe Z Jin, and Michale S Fee. Support for a synaptic chain model of neuronal sequence generation. Nature, 468(7322):394–399, nov 2010.

[28] Emily L Mackevicius, Andrew H Bahle, Alex H Williams, Shijie Gu, Natalia I Denisenko, Mark S Goldman, and Michale S Fee. Unsupervised discovery of temporal sequences in high-dimensional datasets, with applications to neuroscience. Elife, 8:e38471, 2019.

[29] Ofer Tchernichovski, Fernando Nottebohm, Ching Elizabeth Ho, Bijan Pesaran, and Partha Pratim Mitra. A procedure for an automated measurement of song similarity. Animal behaviour, 59(6):1167–1176, 2000.

[30] Jeffrey Cynx. Experimental determination of a unit of song production in the zebra finch (Taeniopygia guttata). Journal of Comparative Psychology, 104(1):3–10, 1990.

[31] Yael Mandelblat-Cerf and Michale S. Fee. An Automated Procedure for Evaluating Song Imitation. PLoS ONE, 9(5):e96484, may 2014.

[32] O Tchernichovski, P P Mitra, T Lints, and F Nottebohm. Dynamics of the vocal imitation process: how a zebra finch learns its song. Science (New York, N.Y.), 291(5513):2564–2569, mar 2001.

[33] Dina Lipkind and Ofer Tchernichovski. Quantification of developmental birdsong learning from the subsyllabic scale to cultural evolution. Proceedings of the National Academy of Sciences of the United States of America, 108 Suppl:15572–15579, sep 2011.

[34] Dina Lipkind, Gary F Marcus, Douglas K Bemis, Kazutoshi Sasahara, Nori Jacoby, Miki Takahasi, Kenta Suzuki, Olga Feher, Primoz Ravbar, Kazuo Okanoya, and Ofer Tchernichovski. Stepwise acquisition of vocal combinatorial capacity in songbirds and human infants. Nature, 498(7452):104–108, jun 2013.

[35] Emily Lambert Mackevicius and Michale Sean Fee. Building a state space for song learning. Current Opinion in Neurobiology, 49:59–68, 2018.

[36] Kenji Doya and Terrence J Sejnowski. Birdsong vocalization learning. Advances in Neural Information Processing Systems 7, 7:101, 1995.

[37] Ila R Fiete, Michale S Fee, and H Sebastian Seung. Model of birdsong learning based on gradient estimation by dynamic perturbation of neural conductances. Journal of neurophysiology, 98(4):2038–2057, oct 2007.

[38] M S Fee and J H Goldberg. A hypothesis for basal ganglia-dependent reinforcement learning in the songbird. Neuroscience, 198:152–70, ec 2011.

[39] Michael S Brainard and Allison J Doupe. Translating birdsong: song-birds as a model for basic and applied medical research. Annual review of neuroscience, 36:489–517, jul 2013.

[40] Nicolas Giret, Joergen Kornfeld, Surya Ganguli, and Richard H R Hahn-loser. Evidence for a causal inverse model in an avian cortico-basal ganglia circuit. Proceedings of the National Academy of Sciences of the United States of America, 111(16):6063–8, apr 2014.

[41] A Hanuschkin, S Ganguli, and R H R Hahnloser. A Hebbian learning rule gives rise to mirror neurons and links them to control theoretic inverse models. Frontiers in neural circuits, 7:106, jan 2013.

[42] Richard Hahnloser and Surya Ganguli. Vocal Learning with Inverse Models. In Principles of Neural Coding, pages 547–564. CRC Press, may 2013.

[43] Matthew D. Golub, Patrick T. Sadtler, Emily R. Oby, Kristin M. Quick, Stephen I. Ryu, Elizabeth C. Tyler-Kabara, Aaron P. Batista, Steven M. Chase, and Byron M. Yu. Learning by neural reassociation. Nature Neuroscience, 21(4):607–616, apr 2018.

[44] Vincent Villette, Arnaud Malvache, Thomas Tressard, Nathalie Dupuy, and Rosa Cossart. Internally recurring hippocampal sequences as a population template of spatiotemporal information. Neuron, 88(2):357–366, 2015.

[45] Usman Farooq, Jeremie Sibille, Kefei Liu, and George Dragoi. Strength-ened temporal coordination within pre-existing sequential cell assemblies supports trajectory replay. Neuron, 103(4):719–733, 2019.

[46] Sam McKenzie, Roman Huszár, Daniel F English, Kanghwan Kim, Fletcher Christensen, Euisik Yoon, and György Buzsáki. Preexisting hippocampal network dynamics constrain optogenetically induced place fields. Neuron, 109(6):1040–1054, 2021.

[47] George Dragoi. Cell assemblies, sequences and temporal coding in the hippocampus. Current opinion in neurobiology, 64:111–118, 2020.

[48] Sam McKenzie, Roman Huszár, Daniel F English, Kanghwan Kim, Fletcher Christensen, Euisik Yoon, and György Buzsáki. Preexisting hippocampal network dynamics constrain optogenetically induced place fields. Neuron, 109(6):1040–1054, 2021.

[49] Marc F Schmidt and Franz Goller. Breathtaking songs: coordinating the neural circuits for breathing and singing. Physiology, 31(6):442–451, 2016.

[50] Kosuke Hamaguchi, Masashi Tanaka, and Richard Mooney. A Distributed Recurrent Network Contributes to Temporally Precise Vocalizations. Neuron, 91(3):680–693, aug 2016.

[51] Emily L Mackevicius, Michael TL Happ, and Michale S Fee. An avian cortical circuit for chunking tutor song syllables into simple vocal-motor units. Nature communications, 11(1):1–16, 2020.

[52] Pengcheng Zhou, Shanna L Resendez, Jose Rodriguez-Romaguera, Jessica C Jimenez, Shay Q Neufeld, Andrea Giovannucci, Johannes Friedrich, Eftychios A Pnevmatikakis, Garret D Stuber, Rene Hen, Mazen A Kheirbek, Bernardo L Sabatini, Robert E Kass, and Liam Paninski. Efficient and accurate extraction of in vivo calcium signals from microendoscopic video data. eLife, 7:e28728, feb 2018.

[53] P. Mitra and H. Bokil. Observed Brain Dynamics. Oxford University Press, USA, 2007.

[54] Tsai-Wen Chen, Trevor J Wardill, Yi Sun, Stefan R Pulver, Sabine L Renninger, Amy Baohan, Eric R Schreiter, Rex A Kerr, Michael B Orger, Vivek Jayaraman, Loren L Looger, Karel Svoboda, and Douglas S Kim. Ultrasensitive fluorescent proteins for imaging neuronal activity. Nature, 499(7458):295–300, jul 2013.

[55] Tatsuo S Okubo, Emily L Mackevicius, and Michale S Fee. In vivo recording of single-unit activity during singing in zebra finches. Cold Spring Harbor protocols, 2014(12):1273–83, ec 2014.

[56] William A Liberti, Jeffrey E Markowitz, L Nathan Perkins, Derek C Liberti, Daniel P Leman, Grigori Guitchounts, Tarciso Velho, Darrell N Kotton, Carlos Lois, and Timothy J Gardner. Unstable neurons underlie a stable learned behavior. Nature Neuroscience, 19(12):1665–1671, ec 2016.

[57] Liron Sheintuch, Alon Rubin, Noa Brande-Eilat, Nitzan Geva, Noa Sadeh, Or Pinchasof, and Yaniv Ziv. Tracking the Same Neurons across Multiple Days in Ca2+Imaging Data. Cell Reports, 21(4):1102–1115, 2017.

